# Complement component C4 regulates amyloid pathology and glial reactivity in mouse model of Alzheimer’s Disease

**DOI:** 10.64898/2026.09.09.750237

**Authors:** Collin J. Nadarajah, Michelle Y. Li, Sohui Park, Jennifer H. Lawrence, Ashish Sharma, Emma P. Danhash, Celeste M. Karch, Erik S. Musiek

## Abstract

The complement system, a proteolytic cascade crucial for innate immune system function, is dysregulated in brain aging and neurodegenerative diseases, including Alzheimer Disease (AD). In models of AD, complement proteins, including C1q and C3, are upregulated and mediate microglial clearance of amyloid plaques and pruning of synapses. Among complement genes, *C4b*, which in mice encodes complement protein C4, is the most highly upregulated in astrocytes in the setting of aging and amyloid pathology. While C4 plays a key role in the complement cascade, there have been no investigations of the direct role of C4 in regulating AD pathology. To probe the function of C4 in amyloid plaque-related pathology, we crossed a germ line *C4b* knockout mouse (C4 KO) to the 5XFAD mouse model of AD-related amyloidosis. To our surprise, we observed striking reductions in amyloid plaque pathology across multiple brain regions in 5XFAD-C4 KO mice relative to standard 5XFAD controls. This reduction in plaque burden stands in sharp contrast to previous reports of C3 deletion in AD models, which increases plaques. Additionally, we observed a reduction in neuroinflammation and peri-plaque glial clustering in 5XFAD-C4 KO mice, suggestive of a role for C4 in regulating overall neuroinflammatory tone in AD. Finally, we observed a phenotypic shift of the peri-plaque microglia to a more reactive disease-associated microglia (DAM) phenotype, indicating that C4 could be an important factor in regulating microglial reactivity in AD. Altogether, our results demonstrate that C4 may have functions beyond the classical complement cascade and may serve as a key facilitator of AD-related glial function and pathology and a possible target for therapeutic modification.

## Introduction

Alzheimer’s Disease (AD) is the most common age-related neurodegenerative disease and the leading cause of dementia in older adults.^1^ Clinical pathological hallmarks of AD include accumulation of neurotoxic extracellular amyloid-beta (Aβ) plaques and intracellular neurofibrillary tau tangles, with the resultant synaptic and neuronal loss leading to progressive cognitive decline.^1,2^ Another pathological hallmark is increased neuroinflammation^3^, manifested by increased glial reactivity of astrocytes^4,5^ and microglia, which contribute to disease pathogenesis.^6,7^ Understanding the neuroinflammatory milieu in AD could reveal new insights into AD pathogenesis and potentially facilitate future therapeutic development. A key component of neuroinflammation is the upregulation and activation of the complement system, a component of the innate immune system that mediates host defense against microbial pathogens in the periphery.^8^ The complement system is comprised of three distinct proteolytic cascades that activate proteins to opsonize targets for phagocytosis, directly lyse cells, and chemoattract immune cells to inflamed tissue.^8^ Complement component 4 (C4) is a key factor in the primary classical complement pathway and serves as one of the principal upstream regulators of complement component 3 (C3) activation, the central confluence of the three complement pathways.^8^ In the central nervous system (CNS), C4 and C3 mediate synaptic engulfment and pruning to refine neural circuits in early development.^9–12^ However, in Alzheimer’s Disease, complement takes on additional functions. Activated complement components such as C3b and C4d are bound to Aβ plaques^13,14^ and transcripts encoding C3 and C4 are highly upregulated in disease-associated glia both in animal models and in humans.^15,16^ Therefore, targeting complement is thought to modulate AD-related neuroinflammation and thereby regulate pathology. Genetic deletion of C3 or inhibiting C3 activation worsened Aβ plaque accumulation in aged AD-mouse models^17,18^; presumably by limiting C3-mediated opsonization and glial clearance of amyloid. Conversely, C3 deletion also partially mitigated the disease-associated microglial synaptic pruning.^16,19^ This dichotomy illustrates that C3 deletion can exert potentially protective effects on plaques, but degenerative effects on synapses, both by altering glial phagocytosis.

While the role of C3 in neurodegenerative diseases has been extensively studies, the function of C4 in neurological diseases remains less defined. In mice, C4 protein is encoded by the *C4b* gene, expression of which is highly upregulated across brain regions during aging, especially within astrocytes.^20,21^ C4 protein is elevated in the CSF of patients with schizophrenia^22^ and increased in the brains of patients with autism-spectrum disorder.^23^ In humans, C4 is encoded by the *C4A* and *C4B* genes located in the immunological MHCIII locus.^24^ Due to the high degree of variability in the MHCIII loci, there are copy number variations (CNV) of the *C4A* and *C4B* genes; increasing gene copy number of *C4A* and *C4B* increases *C4* mRNA expression and positively correlates with schizophrenia risk.^10^ Furthermore, overexpressing mouse *C4b* or human *C4A* in mice leads to synaptic and social behavioral deficits similar to those observed in schizophrenia.^10,25,26^ Conversely, knockdown of *C4b* was shown to improve synaptic function in aged mice.^27^ In neurodegenerative disease, the understanding of C4 function is more limited. C4 protein in the cerebrospinal fluid (CSF) of AD patients positively correlates with increasing AD pathological biomarkers CSF Aβ_42_, phosphorylated tau, and total tau.^28^ A higher proportion of AD patients have high *C4A* and *C4B* copy numbers than healthy controls,^29^ suggesting a possible role that C4 plays in the AD pathogenesis, though the directionality and nature of the relationship is not understood. In AD, transcripts encoding C4 are highly upregulated in glia, including astrocytes^4,15^ and oligodendrocytes,^15,30^ suggesting a role in modulating the neuroinflammatory response in AD.

However, the impact of C4 expression on amyloid plaque pathology and resulting glial and neuroinflammatory responses is unknown. This study sought to directly examine the role of C4 in regulating Aβ pathology and glial responses in the 5XFAD mouse AD model. Our results reveal that *C4b* deletion surprisingly mitigates amyloid plaque deposition, the opposite of what has been reported with C3 deletion. C4b deletion also alters glial responses to plaques, suggesting that C4 protein may exert functions in the brain beyond activation of C3 in the classical complement cascade.

## Results

### C4 deletion ameliorates amyloid-beta plaque pathology

To examine the role of complement C4 in amyloid plaque accumulation and Aβ-related glial responses, we employed a well-described whole-body germ-line *C4b* knockout (*C4b*^*-/-*^) mouse^31^. Mouse *C4b* is the combined murine ortholog of the human genes *C4A* and *C4B*^10,24^, and studying the role of murine *C4b* allows us to investigate potential roles that C4A/C4B play in human disease. We crossed this *C4b* knockout (C4 KO) mouse to the 5XFAD model of AD-related amyloidosis, a common transgenic mouse line which expresses human *APP* and *PSEN1* genes harboring 5 familial AD mutations and develops amyloid-beta (Aβ) plaques throughout the cerebral cortex and hippocampus starting at around 2 months of age.^32^ We generated C4 wild-type (C4 WT) and C4 knockout (C4 KO), 5XFAD positive (5XFAD+) mice. To account for the known sex differences in pathology in the 5XFAD model^32^, we powered experiments to examine each separately. Additionally, we generated C4 WT and C4 KO 5XFAD negative (5XFAD-) as reference non-pathological controls. We aged all mice to 5 months old, an age with moderate plaque accumulation and glial activation, before mice were perfused for analysis **(Fig. 1a)**. We first examined *C4b* transcript levels in the cortex from our mice, and observed increased *C4b* expression in the C4 WT 5XFAD+ mice relative to their C4 WT 5XFAD-littermate controls, aligning with previous findings of *C4b* upregulation in the context of AD.^15,33^ *C4b* mRNA was absent in the C4 KO 5XFAD+ group, confirming deletion **(Supp. Fig. 1a)**.

**Fig. 1:**
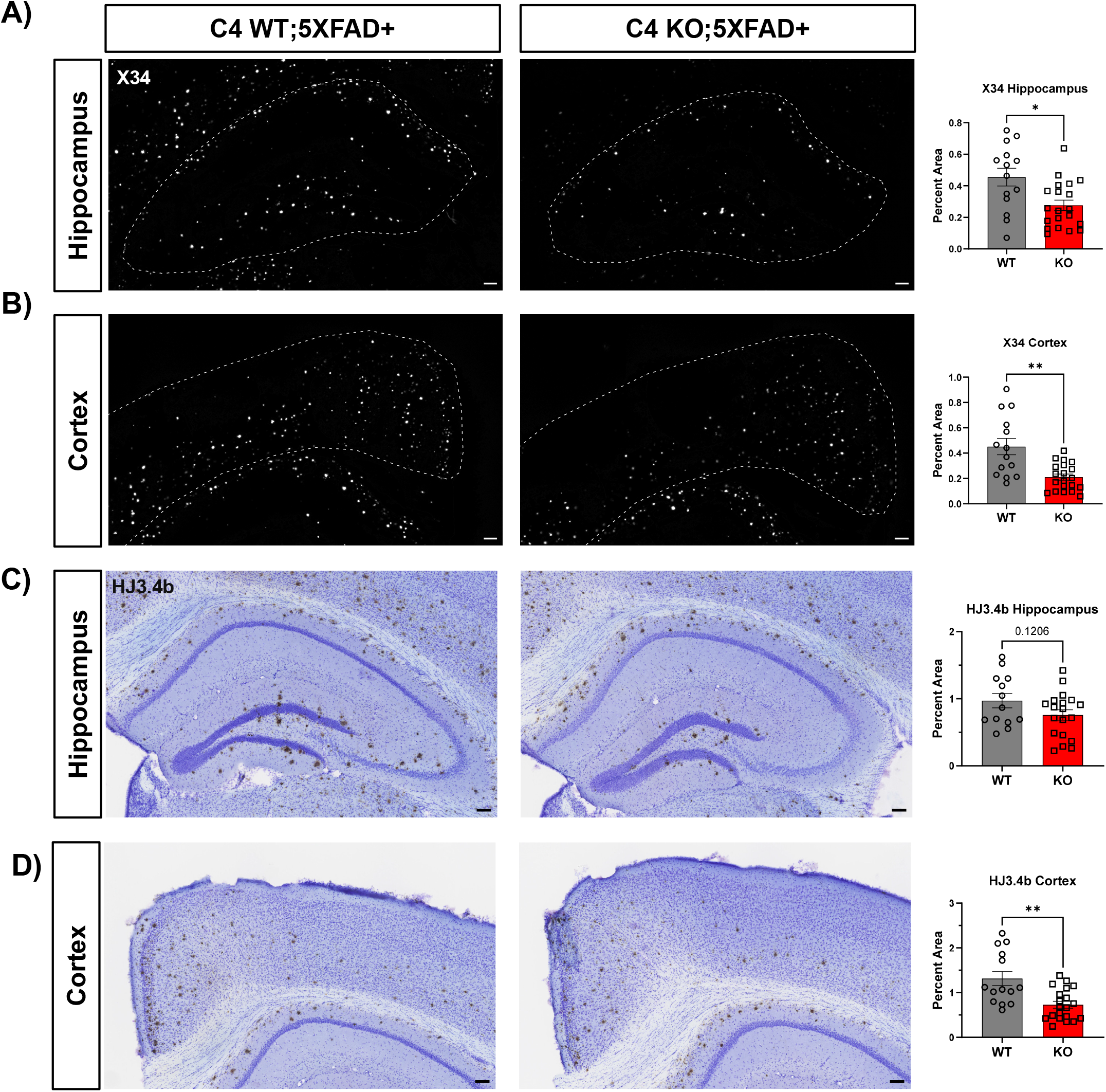
C4 deletion reduces fibrillar and total amyloid-β plaque pathology in multiple brain regions. Fibrillar X34+ amyloid beta (Aβ) pathology is reduced in the **(A)** hippocampus and **(B**) cortex in C4 knockout (KO);5XFAD mice relative to C4 WT;5XFAD controls. Total immunopositive Aβ plaques as measured by HJ3.4b trended to a decrease in the **(C)** hippocampus and were significantly decreased in the **(D)** cortex in the C4 KO mice relative to C4 WT. Graphs show group mean ± SEM. Unpaired 2-tail t-test with Welch’s correction performed unless group variances were significantly different; in that scenario, Mann-Whitney U-test performed. Scale bars, 100 μm. *p < 0.05, **p < 0.01, ***p < 0.001, ****p < 0.0001.

We next examined amyloid plaque pathology in 5-month old 5XFAD mice. To assay the levels of fibrillar Aβ plaques, we utilized X34, a dye that specifically stains amyloid β-pleated sheet structure. We observed striking reductions in X34+ positive plaques across multiple brain regions in the C4 KO; 5XFAD mice relative to C4 WT;5XFAD mice: a 40% reduction in the hippocampus **(Fig. 1c)**, a 54% reduction in the cortex **(Fig. 1d)**, and a 32% decrease in the thalamus **(Supp. Fig. 1b**). We also measured total (diffuse and fibrillar) amyloid pathology, utilizing HJ3.4b, an antibody against Aβ1-13^34^. We detected significant reductions in immunopositive plaques in multiple brain regions: a 40% reduction in the cortex **(Fig. 1f)** and 28% reduction in the thalamus **(Supp. Fig. 1c)**, and a trend toward decrease in the hippocampus **(Fig. 1e)**. This effect was sex-independent, as both males and females showed a relative decrease in plaques of ∼50%, with no statistical difference between them **(Supp. Fig. 1d)**. Since no sex-dependent effect on plaques was observed, we combined both males and females for subsequent analyses. We also characterized the size distribution of plaques in the cortex between C4 WT;5XFAD and C4 KO;5XFAD. We observed a significant decrease in the number of small (0-50 um^2^) and medium (51-100 um^2^) sized plaques **(Supp. Fig. 1e)**. There was also a trending decrease in larger size plaques in the C4 KO, but this was not statistically significant, possibly due to several mice not having any plaques within these size bins. To address possible changes in expression of amyloid-processing genes, we measured the level of transcript for mouse *App* and *Bace1*, by qPCR, and observed no changes for either gene between C4 WT and C4 KO **(Supp. Fig. 1f)**. Together, these findings suggest that C4 is a critical regulator of Aβ plaque pathogenesis.

### C4 deletion broadly reduces Aβ plaque-induced neuroinflammation and neuritic dystrophy

Neuroinflammation is a pathology of AD marked by robust activation of glia, primarily astrocytes and microglia.^1^ Decreasing the neuroinflammatory response could ameliorate pathology and provide targets for disease modifying therapeutics.^35,36^ Therefore, we measured the pathology-induced glial reactivity in the C4 WT and C4 KO 5XFAD+ mice. Because the pathology differences were most pronounced in the cortex, we analyzed the neuroinflammatory response in this region. We measured cortical expression of GFAP, a common astrocytic reactivity marker. We observed a significant 35% reduction in GFAP expression in the cortex of C4 KO;5XFAD mice relative to C4 WT;5XFAD **(Fig 2a)**. To quantify gross microglial reactivity, the other key driver of neuroinflammation, we first measured the expression of microglial marker IBA1. Similar to the astrocytic response, we observed a 25% reduction in IBA1 expression in the cortex in the C4 KO;5XFAD mice **(Fig 2b)**. Additionally, there was more than 30% reduction in LAMP1 immunoreactivity, a marker of peri-plaque dystrophic neurites that are associated with worsened neuronal function^37^ (**Fig. 2c)**. To further characterize microglial activity, we quantified TMEM119, a marker of homeostatic microglia. We observed no changes in TMEM119 expression in the cortex of C4 KO mice relative to C4 WT **(Fig. 2d)**. To gain further insight into the neuroinflammatory environment, we performed RNA expression analysis on cortical tissue, focusing on AD-related inflammatory markers. In line with our IHC data, we observed a significant reduction in expression in neuroinflammatory transcripts such as *Cd68, Gfap, Ctsb, Trem2, Il1b*, and *Lamp1* in the C4 KO;5XFAD mice relative to C4 WT;5XFAD control group **(Fig. 2e)**. Together, these results demonstrate that C4 deletion reduces pathology-induced neuroinflammation broadly in the brain of 5XFAD mice, thereby ameliorating another pathological hallmark of AD

**Fig. 2:**
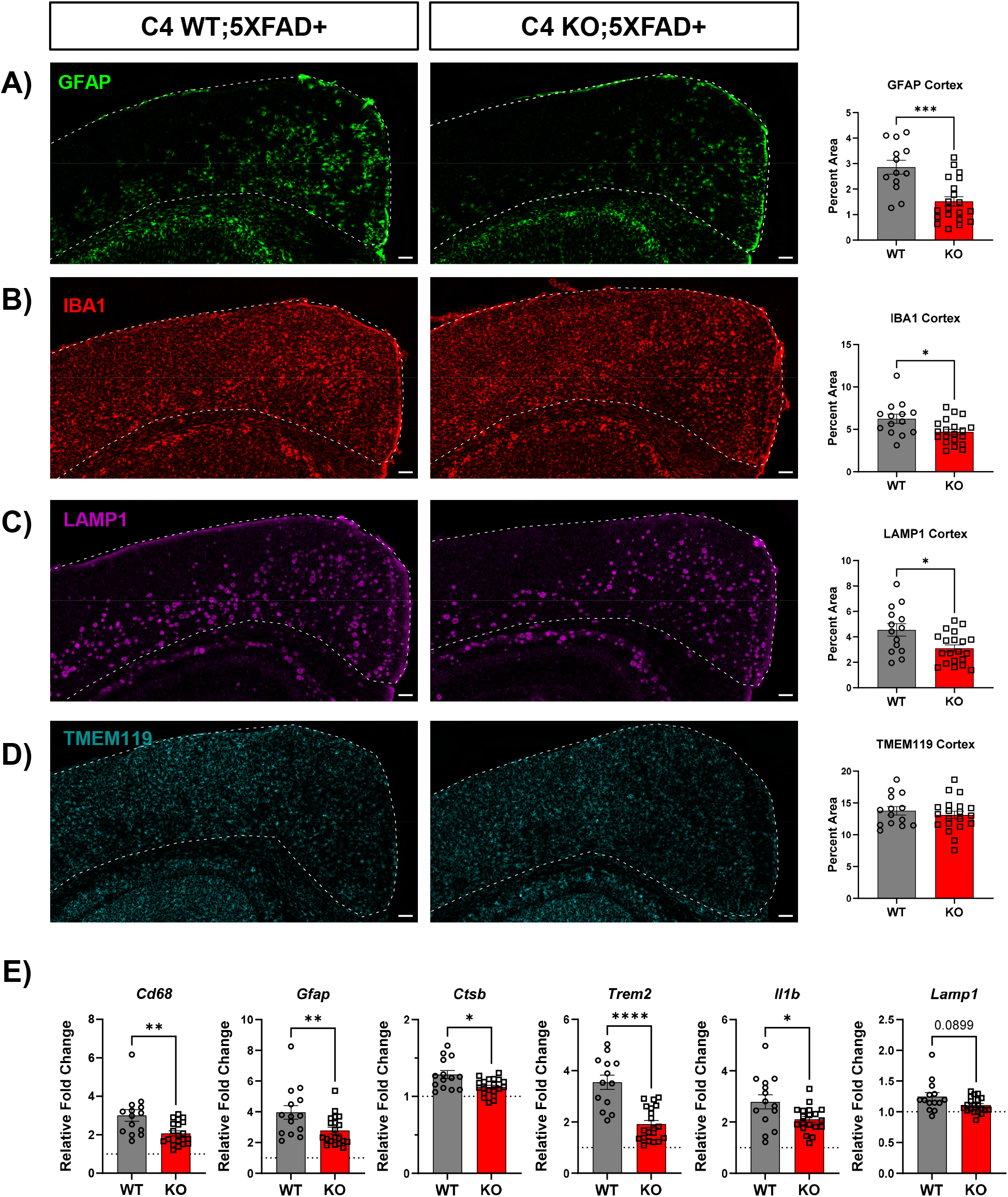
C4 deletion reduces gross plaque-induced neuroinflammation in the cortex. **A)** C4 KO;5XFAD mice had reduced reactive astrocyte GFAP+ percent area in the cortex relative to C4 WT;5XFAD **(B)** C4 KO;5XFAD had reduced IBA1+ percent area in the cortex relative to C4 WT;5XFAD **(C)** ;5XFAD showed reduced dystrophic neurite marker LAMP1 in the cortex relative to C4 WT;5XFAD. (D) Microglia homeostatic marker TMEM119 unchanged between C4 KO;5XFAD and C4 WT;5XFAD. **(E)** C4 KO;5XFAD had reduced neuroinflammatory gene expression in the cortex relative to C4 WT. Relative fold change normalized to C4 WT 5XFAD-negative mice (dashed line). Graphs show group mean ± SEM. Unpaired 2-tail t-test with Welch’s correction performed unless group variances were significantly different; in that scenario, Mann-Whitney U-test performed. N=14-20 per genotype. Scale bars, 100 μm. *p < 0.05, **p < 0.01, ***p < 0.001, ****p < 0.0001

### C4 deletion reduces peri-plaque glia clustering but preserves microglial phagocytosis

Though we observed a reduction in overall glial markers GFAP and IBA1 in the cortex of the C4 KO;5XFAD mice, this can be explained by an overall decrease in amyloid plaque burden. Thus, we sought to characterize the peri-plaque glia, which may mediate the observed plaque reduction phenotype, on a per-plaque basis. To measure the peri-plaque astrocytes and microglia, we used confocal imaging and Imaris 3D image reconstruction to quantify the volume of IBA1+ or GFAP+ voxels within a 15 μm extended surface around individual X34-positive plaques. We then averaged multiple plaques from multiple sections for each mouse. We observed a more than 30% reduction in peri-plaque GFAP+ volume in the C4 KO mice relative to C4 WT, indicating reduced peri-plaque astrocyte clustering **(Fig. 3a)**. Additionally, we observed a 36% reduction in peri-plaque IBA1+ volume in the C4 KO mice relative to C4 WT, demonstrating that there was reduced microglial peri-plaque clustering as well **(Fig. 3b)**. This phenotype was not due to the reduction in plaques in the C4 KO mice as values were normalized to the volume of plaque extended surfaces in the image. Interestingly, despite the decrease in microglia clustering, there was no change in the level of CD68, a marker of phagocytic microglia, within peri-plaque microglia (**Supp. Fig. 2**). This would indicate that even though there is reduced clustering of microglia around the plaques, the microglia maintain their phagocytic protein expression. These results suggest that C4 is required for normal astrocyte and microglial clustering around amyloid plaques.

**Fig. 3:**
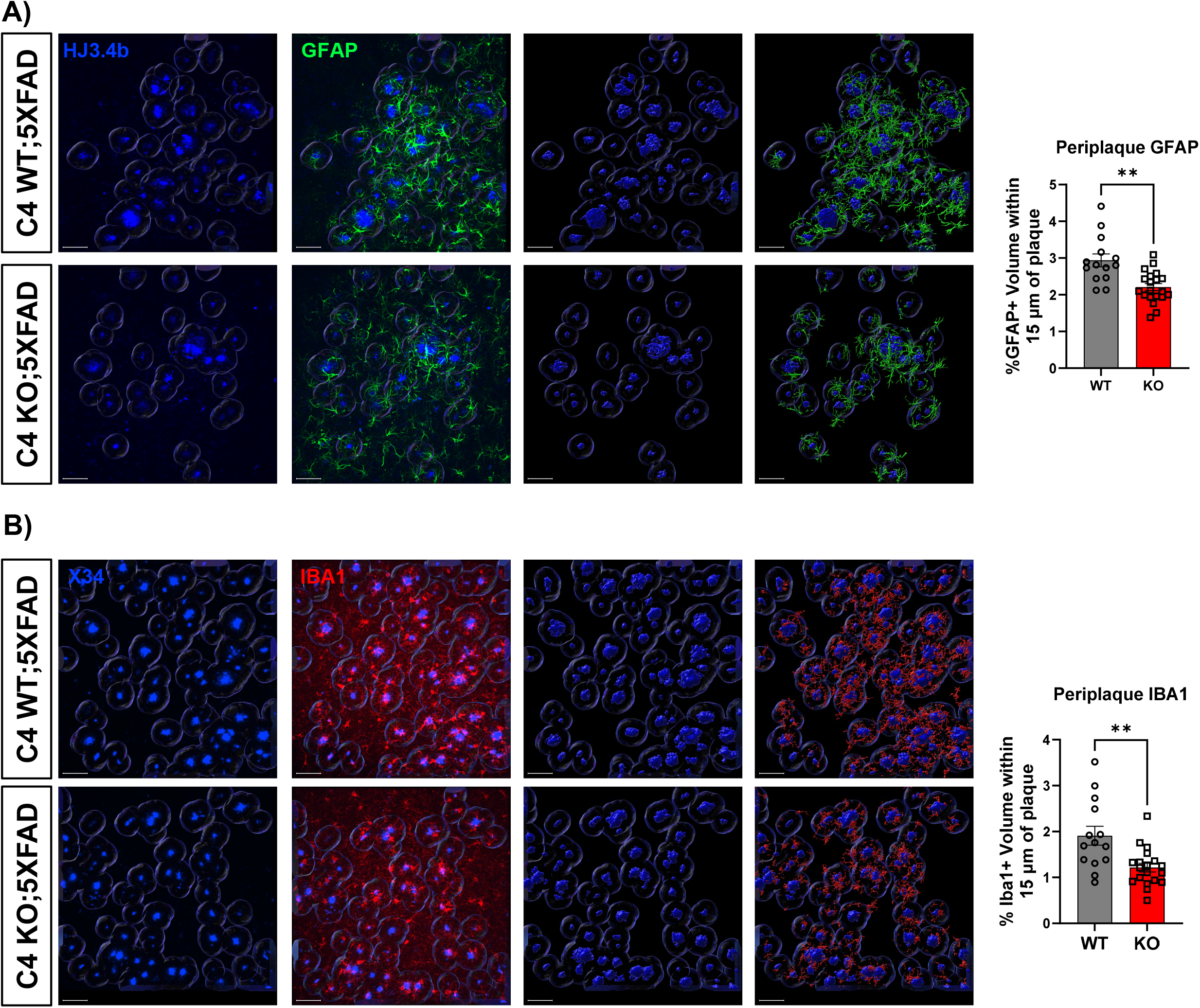
C4 deletion reduces peri-plaque astrocyte and microglial clustering. **(A)** C4 KO reduces GFAP+ astrocyte percent volume within 15 μm of HJ3.4b+ plaques in the cortex in 5XFAD mice. **(B)** C4 KO reduces IBA1+ microglia percent volume within 15 μm of X34+ plaques in the cortex in 5XFAD mice. Panels 1-2 left to right are representative 40x confocal pictures in 3D space. Panels 3-4 are Imaris 3D volume reconstructions of representative images showing analzyed plaque volume, extended plaque surface, and peri-plaque glia. Graphs show group mean ± SEM. Unpaired 2-tail t-test with Welch’s correction performed unless group variances were significantly different; in that scenario, Mann-Whitney U-test performed. N=14-20 per genotype. Scale bars, 50 μm. *p < 0.05, **p < 0.01, ***p < 0.001, ****p < 0.0001

### C4 deletion increases peri-plaque microglial, but not astrocytic, reactivity

Because *C4b* is highly expressed in glia in AD, especially astrocytes^4,15^, it is possible that C4 gates the activation of glia in the context of AD pathology. Though GFAP and IBA1 are general markers of astrocyte and microglia reactivity, respectively, there are more specific markers for AD-specific glial activation. *C3*, in addition to being part of the complement system, is also marker of disease-associated astrocytes (DAA) that is highly increased in neurodegenerative disease.^38^ *Clec7a* is a marker of disease-associated microglia (DAM), a unique microglia transcriptomic and functional signature that is highly upregulated in reactive microglia in AD.^6^ Thus, we quantified levels of C3+ astrocytes and CLEC7A+ microglia, respectively, around plaques, in order to characterize the specific activation state of glia. Surprisingly, there was no change in C3+ GFAP+ positive astrocyte volume within 15 μm of plaques between the C4 WT;5XFAD and C4 KO;5XFAD groups **(Fig. 4)**, suggesting that C4 KO does not change the peri-plaque DAA-like state of astrocytes, even though *C4b* is highly expressed in DAA in humans and mice models. However, there was an 80% increase in CLEC7A+ IBA1+ microglia volume within 15 μm of plaques in the C4 KO;5XFAD mice relative to C4 WT;5XFAD **(Fig. 5)**. This result indicates that C4 could be modulating the shift of homeostatic microglia to a disease-associated phenotype. These results were not affected by the decreased volume of peri-plaque glia as all values were normalized to the amount of peri-plaque glia present in the image. Therefore, our results suggest that C4 can regulate peri-plaque microglia activation state, though the mechanism of this activation remains to be described.

**Fig. 4:**
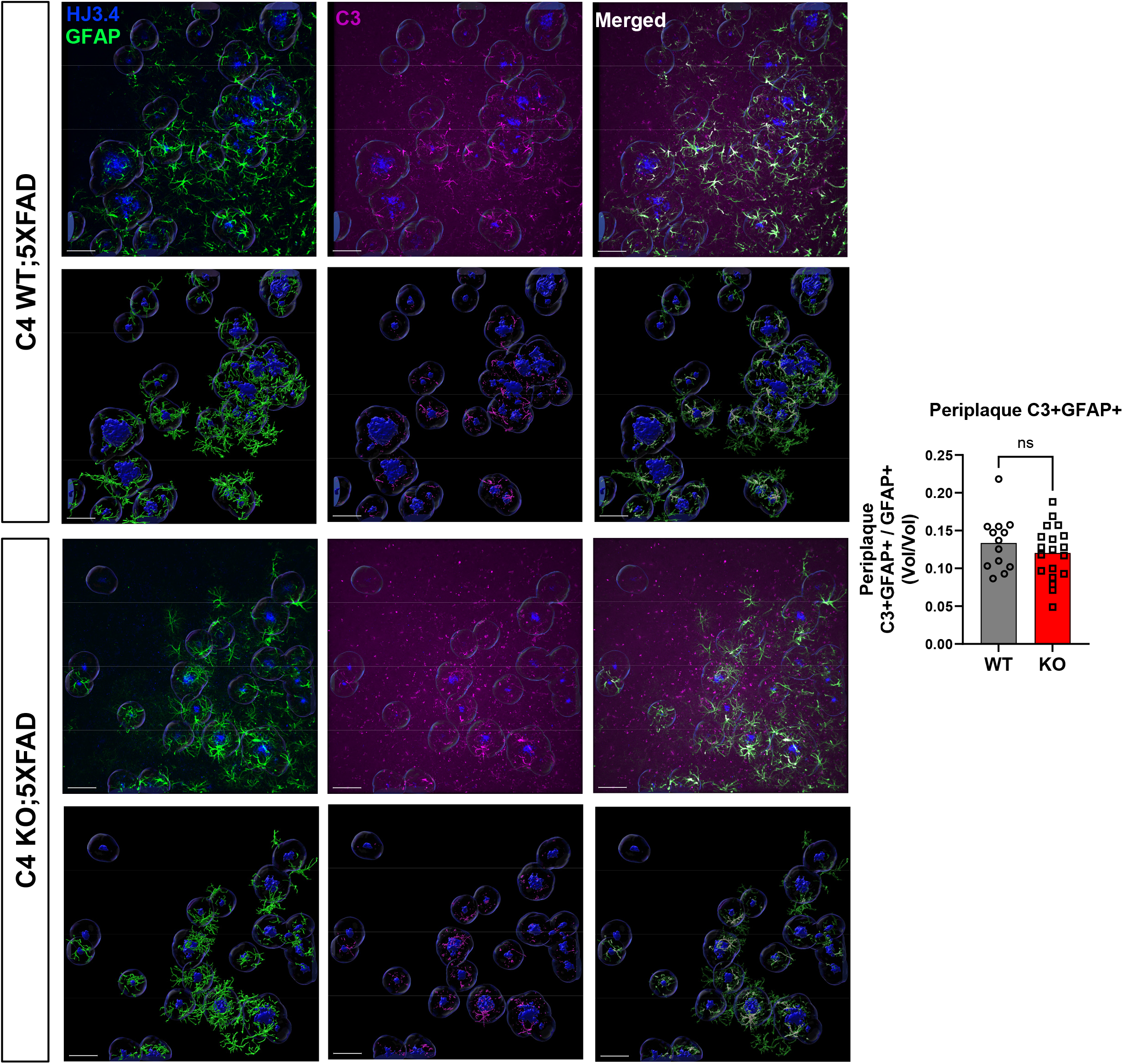
C4 KO does not alter peri-plaque C3 positive astrocyte reactivity in 5XFAD mice. Top row of each genotype is 40x representative confocal image in 3D space showing HJ3.4+ plaques, GFAP+ astrocytes, or C3+ staining with merged image on right panel. Bottom row of each genotype shows Imaris 3D reconstruction of the peri-plaque GFAP, peri-plaque C3, or the colocalized volumes that were analyzed within the 15 μm extended surface. Graphs show group mean ± SEM. Unpaired 2-tail t-test with Welch’s correction performed unless group variances were significantly different; in that scenario, Mann-Whitney U-test performed. N=14-20 per genotype. Scale bars, 50 μm. *p < 0.05, **p < 0.01, ***p < 0.001, ****p < 0.0001

**Fig. 5:**
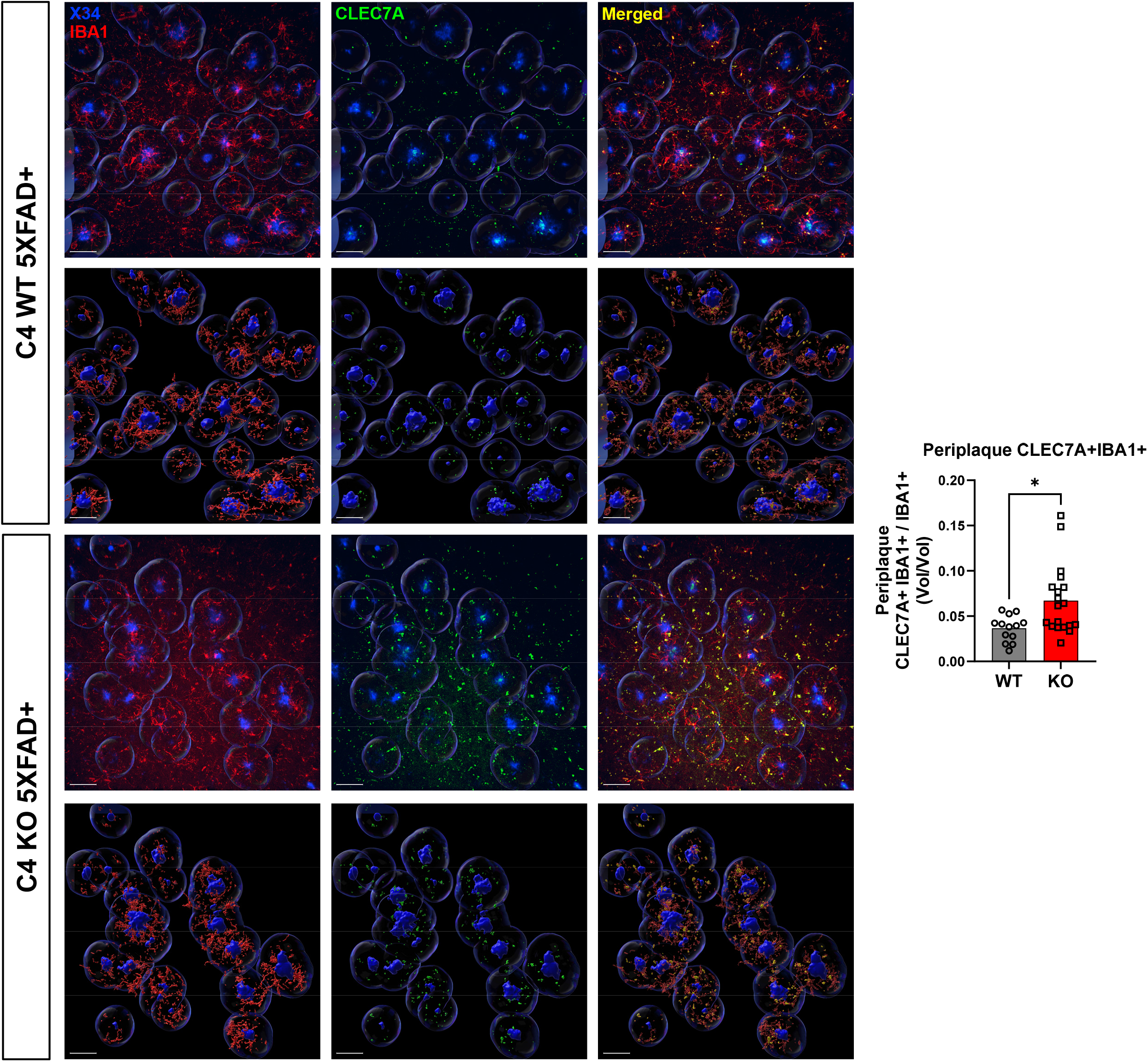
C4 KO increases peri-plaque CLEC7A+ microglia in 5XFAD mice. C4 KO increased the relative percentage CLEC7A in peri-plaque IBA1+ microglia within 15 μm of X34+ positive plaque in 5XFAD mice. Top row of each genotype is 40x representative confocal image in 3D space showing X34+ plaques, IBA1+ microglia, or CLEC7A+ staining with merged image on right panel. Bottom row of each genotype shows Imaris 3D reconstruction of the peri-plaque IBA1, peri-plaque CLEC7A, or the colocalized volumes that were analyzed within the 15 μm extended surface. Graphs show group mean ± SEM. Unpaired 2-tail t-test with Welch’s correction performed unless group variances were significantly different; in that scenario, Mann-Whitney U-test performed. N=14-20 per genotype. Scale bars, 50 μm. *p < 0.05, **p < 0.01, ***p < 0.001, ****p < 0.0001

## Discussion

Complement has long been known to be involved in Alzheimer’s Disease (AD) as activated complement proteins, including C4b and C3b, are found in AD patient brains coating amyloid plaques as part of the overall neuroinflammatory overactivation.^14^ Initially complement proteins were viewed as opsonins targeting plaques for phagocytosis by microglia and astrocytes, but aberrant overactivation targeted neuronal tissue for destruction as well.^39^ Accordingly, disrupting the complement cascade via C3 deletion worsens amyloid plaque burden^17^ but also ameliorates synaptic loss induced by pathology.^19^ Since C4 is critical for C3 activation, the expectation was that *C4b* deletion would mimic prior C3 deletion studies and exacerbate plaque accumulation. However, contrary to the results from C1q^33^ or C3 deletion studies, we have observed here that C4 deletion significantly reduced amyloid beta plaque pathology in 5xXFAD mice. This result is surprising given that C4 is downstream of C1q and upstream of C3 in the classical complement cascade, so one would predict deletion of C1q would lower active C4 levels and that lower C4 levels would result in lower C3 levels. Therefore, the 3 knockout models theoretically should produce similar results. Curiously, C3 levels were increased in the C1q knockout amyloid mouse, which the authors attributed to an increase in activity of the alternative complement pathway.^33^ The alternative pathway is C1q- and C4-independent and involves the auto-hydrolysis of C3 to form active C3b to further activate the downstream complement factors.^8^ It is possible that the alternative pathway activation is neuroprotective and that deletion of C4 abrogates the classical pathway and leads to a compensatory upregulation of alternative pathway activity. Thus, our results suggest a mechanism that occurs independent of the classical complement pathway.

We observed significant reductions in astrocyte and microglial peri-plaque clustering, as well as increased peri-plaque microglial CLEC7A expression in C4 KO;5XFAD mice. In addition to the role complement proteins role as opsonins, they can also function as signaling molecules that can regulate the function of nearby cells. Upon neuroinflammatory activation in AD, astrocytes upregulate C3 and its active cleaved version, C3a, and this C3a is secreted to nearby microglia to regulate phagocytosis.^40^ Microglia recognize C3a through their cognate receptor C3aR, and C3aR inhibition reduced amyloid plaque burden in an amyloidosis model.^40^ C3-C3aR signaling was also an important mediator of tau pathology as genetic deletion of C3aR substantially reduced accumulated tau pathology and resulting neurodegeneration.^41^ C4 can also be cleaved into an active component C4a^8^; C4a is also highly upregulated in patients with schizophrenia and was shown to mediate blood brain barrier permeability in an *in vitro* model.^42^ These results suggest that C4 could also function as a signaling molecule in AD because of the similar upregulation of C4. C4b, a cleavage product of C4, can bind to complement receptor 1 (CR1) in humans and the murine ortholog CR2.^43^ CR1 is expressed on astrocytes in the human brain in AD^44^ and is a known genetic AD risk factor^45^. Additionally, there are single nucleotide polymorphisms (SNPs) in the *CR1* gene that result in CR1 isoforms with increased binding sites for C4b and C3b; these SNPs are further associated with increased AD risk.^46^ Therefore, it is possible that C4 signals through CR1 and that reduction of activity of this signaling axis could ameliorate pathology as observed in our C4 knockout mice. Overactivity of this axis may in part explain why patients with increased gene copy number of C4A, and thus higher C4 expression, have a higher incidence of AD.^29^ However, whether this signaling mechanism is altered in the C4 knockout remains to be defined.

We observed decreases in astrocyte and microglial peri-plaque clustering in the C4 knockout mice relative to control. Other groups have noted reduced astrocytic and microglial peri-plaque clustering in C3 KO relative to C3 WT^19^, a similar phenomenon as observed in our C4 KO mice. This result seems paradoxical as one would expect a closer association of glia to help facilitate clearing of the plaques. It is possible that there is a subtle, yet currently undefined, functional shift in the glia that compensates for the reduced localization. We did not detect a change in C3 positive astrocytes around the plaques, which are thought to be a more reactive neurotoxic subset of astrocytes.^38,47^ This was somewhat surprising given that C4 is highly expressed by astrocyte subpopulations in AD, so loss of C4 altering astrocyte localization but not reactivity suggest different mechanisms are responsible for gating those phenotypes. It is possible that the signaling from reactive microglia are sufficient to induce the reactive astrocyte phenotype^48^, even after the loss of C4. Interestingly, we observed a striking increase in CLEC7A+ microglia around the plaques, indicating an increased switch to the disease-associated microglia (DAM) signature. This reactive microglia population is thought to be a compensatory mechanism that limits neurodegeneration,^6^ and increasing the effectiveness of this select population may prove beneficial. Supporting this idea, C3 deletion decreased general microglial reactivity as measured by phagocytic marker CD68, but other more specific DAM signature genes were not assayed.^17,19^ These C3 KO mice had increased plaque burden, potentially due to a diminished microglial response. Therefore, microglia may be the ultimate mediators of this phenotype

Our study also has limitations as we are utilizing a germline, whole-body deletion of C4 and thus there may be developmental effects that result in the observed plaque reduction. Additionally, because the knockout was in all cell types both in the brain and in the periphery, microglia may not be ultimate effectors of the plaque reduction, and the observed increased DAM reactivity is a byproduct of another unaccounted-for process. Intracellular complement signaling has been implicated in maintaining proper T-cell activation and function^49^, and T-cells have been shown to be critical in driving AD pathology.^50,51^ Therefore the precise mechanism of the plaque reduction remains to elucidated. Cell type specific *C4b* deletion would be needed to further dissect the C4 functions in AD models. Despite these limitations, our study clearly demonstrates that C4 has differing functions than C3 in Alzheimer’s Disease, and that specific study and targeting of each component may be required to generate a comprehensive understanding of complement in neurodegenerative diseases.

## Methods

### Mice

All mice experiments were conducted in accordance with the protocols approved by Washington University in St. Louis Division of Comparative Medicine. 5XFAD hemizygous mice on a C57BL/6J background were from Jackson Labs via the MMRC (strain B6.Cg Tg(APPSwFlLon,PSEN1*M146L*L286V)6799Vas/Mmjax). The 5XFAD transgene was always maintained on the male breeder to prevent variation in plaque burden due to imprinting. Whole-body germline C4 knockout (*C4b*^−/−^) mice were obtained from Dr. John Atkinson (Washington University in St. Louis, St. Louis, MO). All mice were bred at Washington University in St. Louis and maintained on a C57BL/6 background. C4^−/−^ female mice were crossed with one male 5XFAD mouse, yielding a line of C4^+/−^;5XFAD+ mice and a line of C4^+/−^;5XFAD-mice. One male C4^+/−^;5XFAD+ mouse was crossed with C4^+/−^;5XFAD-female littermates, yielding C4^+/+^;5XFAD+, C4^−/−^;5XFAD+, C4^+/+^;5XFAD-, and C4^−/−^;5XFAD-mice. One C4^+/+^;5XFAD+ male was bred with C4^+/+^;5XFAD-female littermates; in parallel, one C4^−/−^;5XFAD+ male was bred with C4^−/−^;5XFAD-female littermates. A cohort of 14 C4^+/+^;5XFAD+ (referred to as C4WT) mice (6 males, 8 females) and 20 C4^-/-^;5XFAD+ (referred to as C4KO) mice (10 males, 10 females) was used for all experiments. All mice were maintained under standard 12h:12h light dark cycles throughout their lives, housed in cages containing 2-5 mice, and sacrificed at 5 months of age.

### Immunohistochemistry

Mice were anesthetized with an intraperitoneal injection of pentobarbital (150 mg/kg) and perfused with ice-cold Dulbecco’s modified PBS (DPBS) containing 3g/L heparin. One hemisphere of each brain was dissected into regions, flash-frozen, and stored at -80°C for other analyses. The other hemisphere was post-fixed in 4% paraformaldehyde (PFA) at 4°C for 48 hours, then cryoprotected in 30% sucrose in 1× phosphate-buffered saline (PBS) at 4°C. 40-micron serial coronal brain sections were harvested on a freezing sliding microtome and stored in cryoprotectant solution (30% ethylene glycol, 15% sucrose, 15% phosphate buffer in ddH20) at -20°C. Two sections, 240 microns apart, in each mouse were stained in all experiments. The following immunohistochemical antibodies were used: X34 (in DMSO, (Sigma-Aldrich, 215294-98-7, 1:2000), HJ3.4 (biotinylated mouse monoclonal, obtained from Dr. David M. Holtzman, H022015, 1:2000), GFAP conjugated to Alexafluor-647 (mouse, Cell Signaling Technology, 3657S, 1:800), C3 (rat, Novus, NB200-540, 1:500), IBA1 (rabbit, Wako, 019-19741, 1:1000), CD68 (rat, BioRad, MCA1957, 1:500), TMEM119 (rabbit, Cell Signaling Technology, 90840S, 1:500), CLEC7A (rat, InvivoGen, mabg-mdect-2, 1:50), LAMP1 (rat, DSHB, 1D4B-s, 1:1000). For sections with fibrillar plaque staining, sections were washed in PBS, incubated in PBSX (PBS + 0.4% Triton-X-100) at RT for 30 min, incubated X34 staining buffer (1:500 10N NaOH added to an aliquot of X34 Wash Buffer) at RT for 20 min, then washed in X34 Wash Buffer (60% PBS, 40% EtOH). Fluorescent antibody co-staining was then done in PBS as described below. For sections with C3 staining, 2 sections per animal were heated in citrate buffer (Invitrogen, 005000) for 30 min at 80°C for antigen retrieval. Fluorescent antibody co-staining was then done in PBS as described below. For sections with glial staining only, sections were washed in 1x Tris-buffered saline (TBS), blocked in TBSX (TBS + 0.4% Triton X-100 (Sigma-Aldrich)) containing 3% (20% for CLEC7A stains) donkey serum at RT for 60 min, and incubated in primary antibodies diluted in TBSX containing 1% (3% for CLEC7A stains) donkey serum overnight at 4°C. Sections were then washed in TBS, incubated in TBSX with 1:1000 donkey fluorescent secondary antibody at RT for 60 min, and mounted on slides using ProLong Gold antifade reagent (Invitrogen, P36934) before coverslipping. For total plaque staining, sections were washed in TBS, incubated in 0.3% hydrogen peroxide in TBS for at RT for 10 min, washed in TBS, blocked in 3% milk in TBS at RT for 30 min, then incubated in 1:1000 HJ3.4B in 1% Milk-TBS-X at 4°C overnight. The next day, sections were washed in TBS, incubated in 1:400 ABC Elite (Vector PK-6100) in TBS at RT for 60 min, washed in TBS, incubated in DAB (Sigma-Aldrich 5905-50TAB) + 100ul 4% Nickel solution (in water) + 20ul 30% hydrogen peroxide for 7 minutes, washed in TBS, then mounted and allowed to dehydrate at RT overnight. Sections were counter-stained with Cresyl violet. Sections were then covered in ProLong Gold antifade reagent (Invitrogen, P36934) and coverslipped.

### Imaging and Image analysis

Epifluorescent microscopy was conducted using a Keyence BZ-X810 microscope. The 10x objective was used to take images of the entire hemi-brain. Exposure settings were determined for each antibody in each experiment following an initial survey of the tissue and applied across all samples. Uncompressed images were stitched on the channel containing the most background signal. 10x images were saved as TIFF files. Brightfield microscopy was conducted on the NanoZoomer digital pathology system to image sections stained with HJ3.4. The 20x objective was used to take images of the entire slide. Images of individual sections were exported as TIFFs using NDP.view2, a digital slide viewer software. Image deconvolution was performed in ImageJ using the Color Deconvolution 2 plugin (Dr. Gabriel Landini, University of Birmingham, Birmingham, England). 10x images of the HJ3.4 color were saved as TIFF files. Epifluorescence and brightfield images were analyzed using ImageJ software (NIH). TIFF files were opened in ImageJ and regions-of-interest (ROIs) were drawn and saved. Images were converted to 8-bit grayscale and the Subtract Background function was applied to reduce noise. Optimal brightness and threshold values were determined by comparison across multiple images with variable staining intensities, then applied to all images in the cohort. Using the Analyze Particles function, percent area of the ROI stained was quantified for each section. Percent area quantifications for two sections per mouse per region were analyzed and averaged. Statistics were then performed on the mouse averages between groups (C4KO and C4WT). Z-stack images were taken on a Nikon AXR laser scanning confocal microscope using the 40x oil immersion objective on the Galvano scanning setting. Averaging, dwell time, laser intensity, and laser gain settings were determined by surveying tissue with varying staining intensities and applied to all samples. Z-stack images were imported and analyzed in the visualization and analysis software Imaris. 3D surfaces of X34+ and HJ3.4+ plaques were rendered, then a surface dilation function was performed to generate a peri-plaque region 15μm from X34+ or HJ3.4+ plaque surfaces. 3D surfaces of glia stained with GFAP, C3, IBA1, CD68, CLEC7A, TMEM119, and were rendered. Peri-plaque clustering was determined by calculating the volume of glial surfaces within the plaque dilated surfaces using a surface colocalization function. Glia volumes were normalized to the dilated surface volume. Peri-plaque glia activation was determined by obtaining the colocalized volume of gross glia markers with activation markers within the dilated surface. Peri-plaque volumes across all plaques in each section were summed, and the sums were averaged between two sections in each mouse. Statistics were performed mouse averages between groups (C4KO and C4WT).

### qPCR

Cortical tissue was added to Eppendorf SafeLock tubes and immersed in 500ul of TRIzol (Thermo Fisher Scientific) and beads (company), then homogenized in a Bullet Blender Homogenizer (Next Advance) at level 8 for 3 minutes. Chloroform was added according to a 1:6 chloroform:TRIzol ratio, samples were vigorously mixed and centrifuged at 12500g for 15 minutes at 4°C, and the supernatant was collected into separate tubes. RNA was extracted using the PureLink RNA Mini Kit as described by the manufacturer’s protocol. RNA concentrations in each sample were determined using a Nanodrop spectrophotometer. Samples of 1000ng RNA per 20 µL were made to synthesize cDNA using an RNA-cDNA reverse transcription kit (Applied Biosystems/Life Technologies). Samples were analyzed via microfluidic qPCR array analysis using a Fluidigm Biomark HD system with TaqMan primers (Life Technologies) at the Washington University Genome Technology Access Center (GTAC).

### Protein Extraction

Cortical tissue was added to Eppendorf SafeLock tubes and immersed in 400ul of RIPA buffer (150mM NaCl, 50mM Tris, 0.5% deoxycholic acid, 1% Triton-X 100, 0.1% SDS, 5mM NaF, and 1mM sodium orthovanadate) with 1% protease & phosphatase inhibitor and Rnase-free beads, homogenized using the Bullet Blender Homogenizer (Next Advance) at level 8 in 30 second intervals, repeating as needed, and centrifuged at 4°C at 10,000g for 5 minutes. The supernatant was collected as the soluble fraction. The remaining pellet was incubated in six parts of 5M guanidine HCL with 50mM TRIS and 1% protease & phosphatase inhibitor. Samples were sonicated at 10% amplitude for 30 seconds in a 4°C water bath sonicator (Qsonica), incubated at RT for 2 hours with agitation, and centrifuged at 4°C for 20 minutes at 15,000g. The supernatant was collected as the insoluble fraction. Pierce BCA Protein Assay (Thermo Fisher Scientific) was performed as described by the manufacturer’s protocol.

### Western Blot

Protein from cortical tissue was extracted as described above. 40μg of protein were added to 40ul Western blot samples were made by dissolving 40ug of protein in Laemlli sample buffer (BioRad) and Milli Q (Millipore) water. Samples were heated at 95°C for 5 minutes and separated using SDS-polyacrylamide gel electrophoresis at 80V for 1 hour, then transferred to a 0.2μm immunoblotting membrane (PVDF) with electrophoresis at 30V for 1.5 hours. The membranes were rinsed 0.05% TBST (TBS + Tween 20), blocked in 8% milk in TBST for an hour at RT, and incubated in primary antibody diluted in TBST overnight at 4°C. The following primary antibodies were used: Aβ (rabbit, Thermo, 51-2700, 1:1000), PSEN1 (rabbit, CST, 5643, 1:1000), BACE1 (rabbit, Abcam, EPR3956, 1:1000), β-actin (rabbit, CST, 4970, 1:8000). The membranes were washed in TBST, then incubated in anti-rabbit horseradish peroxidase (HRP)-conjugated secondary antibody in 1% bovine serum albumin (BSA) for 1 hour at RT. The membranes were briefly dipped in Super Signal Wes Pico PLUS Chemiluminescent Substrate (Thermo Fisher Scientific, 34577) and imaged on a ChemiDoc Imaging System (BioRad). Images were analyzed using ImageJ and brightness was normalized to actin.

### Statistics

Statistical analyses were performed using GraphPad Prism software version 10.4.0, including unpaired two-tailed t-tests (for two groups with equal variance, as determined by F-test), or by 2-tailed T-test with Welch’s correction if variances were unequal. For comparisons with two variables, two-way ANOVA with Tukey’s multiple comparison was used. Outliers were identified using Grubb’s test and excluded if their significance level exceeded alpha = 0.05. Data are presented as the mean ± standard error of the mean (SEM), with significance levels indicated by asterisks. P-values greater than 0.05 were labeled as ‘ns’ or left unlabeled, while those below 0.05 were considered significant (*p < 0.05; **p < 0.01; ***p < 0.001; ****p < 0.0001).

## Acknowledgements

This work was supported by NIH grants R01AG054517 and R01AG063743 (ESM). Confocal imaging and Imaris image analysis was performed at the WashU Center for Cellular Imaging (WUCCI), which is supported by Washington University School of Medicine, The Children’s Discovery Institute of Washington University and St. Louis Children’s Hospital (CDI-CORE-2015-505 and CDI-CORE-2019-813) and the Foundation for Barnes-Jewish Hospital (3770 and 4642).

