## Supplemental Figures 1 & 2 for "Complement component C4 regulates amyloid pathology and glial reactivity in mouse model of Alzheimer’s Disease"

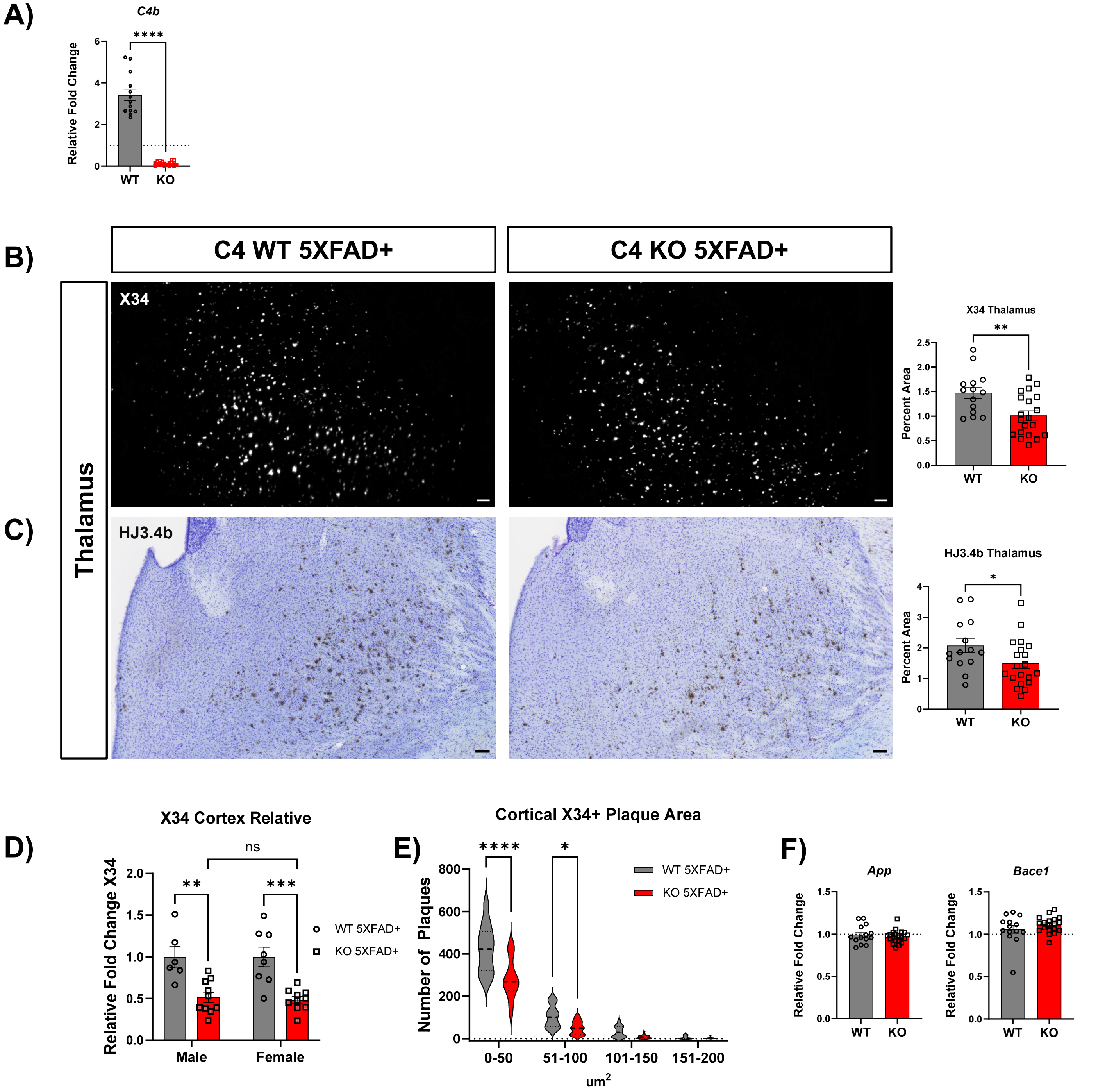


**Supp. Fig. 1: C4 deletion reduces fibrillar and total amyloid-β plaque pathology in multiple brain regions. (A)** *C4b* mRNA levels in cortex samples from C4 WT 5XFAD+ and C4 KO 5XFAD+ mice. Relative fold change normalized to WT control. **(B)** X34+ and **(C)** HJ3.4+ plaques are reduced in the thalamus in C4 KO 5XFAD+ mice relative to C4 WT **(D)** No difference observed in cortical X34+ plaque reduction between males C4 KO and female KO relative to WT mice of the same sex. **(E)** Small and medium sized cortical X34+ plaques are significantly reduced in C4 KO relative to WT. **(F)** No difference observed in *App* and *Bace1* transcript levels in C4 KO relative to WT. Graphs show group mean ± SEM. Unpaired 2-tail t-test with Welch's correction performed unless group variances were significantly different; in that scenario, Mann-Whitney U-test performed. N=14-20 per genotype except for 4.3b, where N= 6-10 per sex per genotype. Scale bars, 100 μm. *p < 0.05, **p < 0.01, ***p < 0.001, ****p < 0.0001.

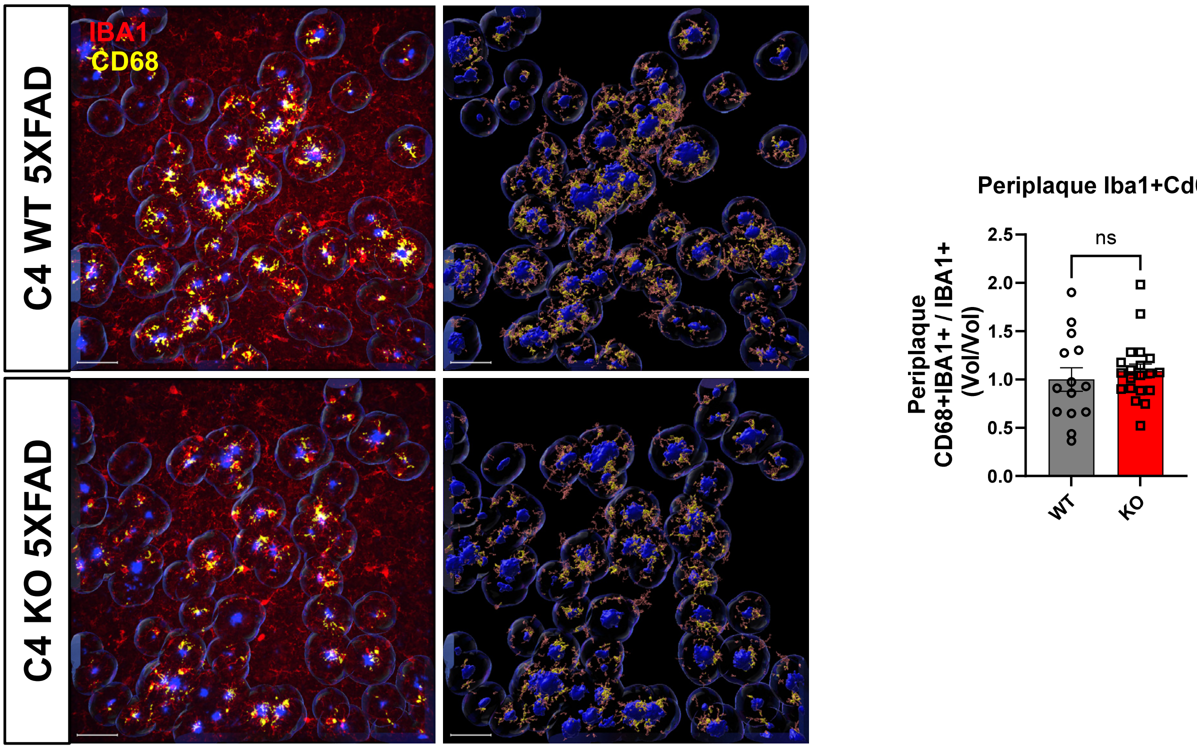


**Supp. Fig. 2: C4 deletion does not alter peri-plaque CD68+ microglia**
No difference between C4 KO and C4WT in relative percentage of CD68-IBA1 colocalization within 15 µm of X34+ plaques. First panel is representative 40x confocal picture, second panel is Imaris 3D reconstruction of representative image. Graphs show group mean ± SEM. Unpaired 2-tail t-test with Welch's correction performed unless group variances were significantly different; in that scenario, Mann-Whitney U-test performed. N=14-20 per genotype. Scale bars, 50 µm. $*p < 0.05$, $**p < 0.01$, $***p < 0.001$, $****p < 0.0001$
